# A novel lineage-tracing mouse model for MmuPV1 infection enables *in vivo* studies in the absence of cytopathic effects

**DOI:** 10.1101/2021.08.19.456940

**Authors:** Vural Yilmaz, Panayiota Louka, Katerina Strati

## Abstract

Human papillomaviruses (HPVs) are DNA viruses that ubiquitously infect humans and have been associated with hyperproliferative lesions. The recently discovered mouse specific papillomavirus, MmuPV1, provides the opportunity to study papillomavirus infections *in vivo* in the context of a common laboratory mouse model (*Mus musculus*). To date, a major challenge in the field has been the lack of tools to identify, observe and characterize individually the papillomavirus hosting cells and also trace the progeny of these cells over time.

Here, we present the successful generation of an *in vivo* lineage-tracing model of MmuPV1-harboring cells and their progeny by means of genetic reporter activation. Following the validation of the system both *in vitro* and *in vivo*, we used it to provide a proof-of-concept of its utility. Using flow-cytometry analysis, we observed increased proliferation dynamics and decreased MHC-I cell surface expression in MmuPV1-treated tissues which could have implications in tissue regenerative capacity and ability to clear the virus. This model is a novel tool to study the biology of the MmuPV1 host-pathogen interactions.

## Introduction

Human papillomaviruses (HPVs) are DNA viruses which are linked to 5% of human cancers (McBride, 2017) and are responsible for commensal infections. Mucosotropic high-risk HPVs, a group which includes HPV16 and HPV18, have been associated with malignancies, mainly cervical cancer, other anogenital cancers and a subset of head and neck squamous cell carcinoma (zur Hausen, 2009). For this reason, the contribution of these viruses to cancer has been extensively studied with significant advances in cancer prevention, prophylaxis, and detection. In addition to representing a significant source of pathogenesis, it is clear that papillomaviruses abundantly colonize human tissues (Hannigan et al., 2015), an aspect of their biology which is more poorly understood.

Since differentiation of stratified epithelia is essential for the productive HPV replication cycle (Roden and Stern, 2018), and due to the species specificity of papillomaviruses, it has been challenging to study the host-virus interaction and the progress of infection of HPV in the laboratory. Furthermore, the outcome of HPV infection is greatly influenced by the host immune system and inflammation induced by tissue damage that is required in order to expose the underlying basal layer to the virus (Amador-Molina et al., 2013). Thus, the best system to study virus-host interaction taking into consideration all the factors that influence infection outcome is to explore this *in vivo*.

This is now possible due to the discovery of a mouse specific papillomavirus, MmuPV1 (Hu et al., 2017). This mouse papillomavirus provides the opportunity to study papillomavirus infections in the context of a small common laboratory animal for which abundant reagents are available and for which many strains exist. The genome of the human and mouse papilloma virus is similar with the only exception of the lack of E5 gene in the mouse virus. The major oncogenes E6 and E7 share similarities in the mouse and human papilloma virus and thus by studying the mouse papilloma virus– host interaction we can gain a better understanding on how the presence of the virus alters the host cell behavior over time (Spurgeon and Lambert, 2020).

Existing approaches in the field have relied on long-term phenotypes leading to clearer understanding regarding the mechanisms of oncogenesis. This remains true in the development of a novel mouse model of spontaneous high-risk HPV E6/E7-expressing carcinoma (Henkle et al., 2021), and two novel models based on infection (Wei et al., 2020) and transmission (Spurgeon and Lambert, 2019). Reliance on cytopathic effects has resulted in a severe understudy of the majority papillomavirus infections which do not lead to overt signs of infection. These infections remain critical to study as they can offer significant insights on the prevention of substantial morbidities such as laryngeal respiratory papillomatosis, genital wart recurrence etc. (Egawa and Doorbar, 2017). This underscores the clear need for a model where one can observe directly the MmusPV1 infected cells and furthermore, trace them over time during the course of the infection.

Here, we have designed and developed a model for *in vivo* lineage tracing of the cells initially harboring MmuPV1. We have created a MmuPV1-lox-Cre-lox plasmid which we deliver to the tail skin of reporter Rosa26-lox-Stop-lox-YFP mice. Expression of self-deleting Cre from the plasmid results in recombination of the *loxp* sites and excision of the sequence flanked by the *loxp* sites. Therefore, introduction of the plasmid in mouse cells leads to the production of Cre recombinase and loss of the Cre sequence and its promoter from the plasmid and more importantly, recircularization of the MmuPV1 genome and the loss of the stop codon upstream of the yellow florescence protein (YFP) gene. This lets us observe and trace longitudinally, cells that initially harbored the plasmid containing the viral genome. Since YFP expression is genetically activated, it is conserved in the following progeny of the initial cell that taken up the plasmid. This model provides, for the first time, a promising tool to study the biology of MmuPV1 infection and its target cells along with the impact of the virus on these cells.

## Materials and Methods

### Plasmids

To generate a plasmid with a self-deleting Cre sequence and the full MmuPV1 genome, we combined sequences from four plasmids: puC19-MmuPV1 (gift from Paul F. Lambert), pCAG-cre (a gift from Connie Cepko (Addgene plasmid # 13775; http://n2t.net/addgene:13775; RRID: Addgene_13775)), small_pMA-T and large_pMA-T (synthesized by Invitrogen). To construct the final plasmid, we first PCR amplified the sequence in large_pMA-T that contains the lox and part of the E6 sequence using primers to introduce PstI and HindIII restriction sites in the 5’ and 3’ ends of the sequence respectively. Using PstI and HindIII we cloned lox-E6 in the pCAG-Cre plasmid. In this new plasmid we subcloned the L1-lox sequence from small_ pMA-T plasmid using blunt end ligation (L1-Lox sequence was isolated using SacI and KpnI restriction enzymes, the Cag-cre plasmid was digested with SalI and T4 DNA polymerase was used to generate blunt ends). We refer to this plasmid as plox-cre-lox, which was sequenced to confirm the correct insertion of L1-lox-Cre-lox-E6 sequence. To generate the plasmid for animal infection, we removed the ampicillin bacterial selection cassette from lox-cre-lox and MmuPV1 plasmids using XhoI and XbaI. The two fragments (lox-cre-lox and MmuPV1) were gel purified and ligated overnight at 16°C (ligation was done using a ratio of 1:2.5, MmuPV1 to lox-cre-lox). Complete plasmid sequences are provided in the supplementary file.

### Tissue culture, Transfection, PCR, RT-PCR

Mouse embryonic fibroblasts (MEFs), isolated from C57BL/6 Rosa26-lox-STOP-lox-YFP pregnant mice, were used for in vitro transfections. MEFs were maintained in Dulbecco’s Modified Eagle Medium (DMEM, GibcoTM, cat. no 41965039) supplemented with 10% v/v fetal bovine serum (FBS, GibcoTM, cat. no 10500064) and 1% v/v penicillin/streptomycin (GibcoTM, cat. no 15070063). In vitro transfections were performed with FuGENE^®^6 Transfection Reagent (Promega, cat. no E2691) as described previously (Hong et al., 2009).

DNA and RNA isolations were performed using TRIZOL (Invitrogen) following the manufacturer’s instructions. We used RT-PCR to confirm expression of Cre (Cre forward primer: 5’ - ACG AGT GAT GAG GTT CGC AA – 3’, Cre reverse primer: 5’ - CGC CGC ATA ACC AGT GAA AC- 3’, GAPDH was used as a housekeeping gene) and PCR to determine lox-recombination and removal of the sequence flanked by loxp sites from the plasmid (Cre loss forward primer: 5’ - ACT GTT TAC TGG GGG CTT AC – 3’ and cre loss reverse primer: 5’ - GGC AGA ACA CAA TGG AAC TC – 3’ ). RT-PCR analysis was performed to assess the presence of viral transcripts from pCAG-Cre and MmuPV1-lox-cre-lox transfected MEFs. The following primer pairs were used and data were analyzed with Rotorgene 6000 Series Software (Qiagen): GAPDH: 5′- ACT CCA CTC ACG GCA AAT TC-3′ and 5′- TCT CCA TGG TGG TGA AGA CA-3′, E1/E4: 5′- GAA CTC TTC CCA CCG ACA CC-3′ and 5′- AAG GTC CTG CAG ATC CCT CA-3′, E6: 5′- CTT TTC AGA GGC AGT AAG GA-3′ and 5′- CTT TTCA GAG GCA GTA AGG A-3′, L1: 5′- TAT ATA ACA TCA TCG GCA AC-3′ and 5′- TGC TTC CCC TCT TCC GTT TT-3′.

### Mice

Rosa26-lox-stop-lox-YFP immunocompetent mice, obtained from the mouse facility of the Cyprus Institute of Neurology and Genetics (CING), were of a pure C57BL/6 genetic background. Wild-type immunocompetent FVB/N mice (obtained from CING) were also used where stated. All mice were used at 6-12 weeks of age and were sex-and age-matched within experiments. All the genotypes were confirmed by means of PCR. Mice were housed at the University of Cyprus, in accordance with regulations and protocols approved by the Cyprus Ministry of Agriculture.

### Ethics statement

This study was carried out in strict accordance with the recommendations in the Guidelines for the Protection of Laboratory Animals of the Republic of Cyprus. The animal facility is licensed by the Veterinary Services (Republic of Cyprus Ministry of Agriculture and Natural Resources), the government body in charge of approving and overseeing laboratory animal work in Cyprus (license number CY.EXP.105) and the protocol was approved by the same authority (License number CY/EXP/PR. L1/2019).

### Plasmid delivery

The back skin of 6-9 weeks old female Rosa26 or FVB/N mice were shaved and a single dose of 1000 mJ/cm2 UVB irradiation was applied for immunosuppression as previously described (Dorfer et al., 2021). Following the UVB radiation, the mice were given at the same day with 2 × 10^9^ DNA copies of MmuPV1-lox-cre-lox or CAG-Cre plasmid in a 10 ul of PBS solution applied at the base of the tail skin by a superficial scarification with a pipette tip to expose the tail basal layer. Tissue samples were then harvested at the specified time points.

### Tissue procurement and Processing

Tail tissue sections from the base of the tail were harvested in a rectangular manner and were cut in half, washed with 1X PBS, fixed in 4% paraformaldehyde (PFA), and embedded in optimal cutting temperature compound (OCT) and frozen on dry ice before storing in −80 °C. Frozen tissues were then sectioned (20 microns thick) using a cryostat and mounted on the positively charged Thermo Scientific™ SuperFrost Plus™ Adhesive slides.

### RNA in situ hybridization

MmuPV1 viral transcripts were detected using RNAscope 2.5 HD Assay-Brown (cat. no. 322300) (Advanced Cell Diagnostics, Newark, CA) according to manufacturer instructions with probes specific for MmuPV1 E6/E7 (Cat #409771) and for MmuPV1 E1/E4 (Cat #473281) as described previously (Spurgeon et al., 2019; Xue et al., 2017). Tissue sections were treated following protease treatment for 30 min at 40°C followed by the probe hybridization. Tissues were then counterstained with hematoxylin and mounted in Cytoseal media (Thermo Fisher Scientific).

### Immunofluorescence

After antigen retrieval was performed in a microwave using 10 mM citrate buffer, the tissue sections were permeabilized and blocked at room temperature for 1 hour with a blocking buffer containing 0.5% skim milk powder, 0.25% fish skin gelatin and 0.5% Triton X-100. Then, sections were washed with PBS, and stained with purified primary antibodies in blocking buffer at 4°C overnight. Tissues were then washed with PBS three times, stained with secondary antibodies at room temperature for one hour, counterstained with Hoechst Dye, mounted in Prolong mounting media (Prolong Gold Antifade reagent, Invitrogen, cat. no. P36930) and sealed with nail polish.

The following antibodies were used for detecting mouse antigens by immunofluorescent staining: rabbit-anti-K14 (polyclonal, 1:1000, Covance, PRB-155b), goat-anti-GFP (polyclonal, 1:100, SICGEN, AB0020-200), Alexa Fluor® 488 donkey-anti-goat (polyclonal, 1:250, Jackson, 705-545-147), Alexa Fluor® 488 donkey-anti-mouse (polyclonal, 1:250, Jackson, 715-545-150), FITC donkey-anti-rabbit (polyclonal, 1:250, Jackson, 711-095-152), Alexa Fluor® 594 donkey-anti-rabbit (polyclonal, 1:250, Jackson, 711-585-152).

### Flow cytometry

After mice were sacrificed and tail skins were extracted as described above, the tail samples were incubated in a 0.25% trypsin containing EDTA solution (Sigma-Aldrich) overnight at 4°C. The next day, epidermis was separated from dermis by using tweezers, the tissue was cut into small pieces and was mechanically dissociated by using a handheld homogenizer (POLYTRON^®^ PT 1200 E). Subsequently, the resulting single cell suspensions were washed with PBS and cells were incubated with fluorescently conjugated antibodies for 30 minutes at 4°C. After surface marker stainings, cells were fixed and permeabilized for 10 min with 2× BD lysing solution (BD Biosciences) + 0.1% Tween, washed, and then stained in PBS for intracellular Ki-67 staining. The following antibodies (indicating target-fluorochrome) were purchased from Biolegend and were used according to the manufacturer’s instructions: MHC-I(H-2Kb)-Brilliant violet 421, Ki-67-PerCP/Cy5.5 and TLR9-PE. Multi-parametric flow cytometric analysis was performed using a Bio-Rad S3e Cell Sorter flow cytometer and analyzed using FlowJo software (Treestar). The population of live cells was detected depending on their size and complexity (FSC-SSC gate) and later the doublets were excluded (FSC-Height/FSC-Area gate).

### Microscopy

For the visualization of tissue sections for RNAscope analysis, the Zeiss Axio Observer.A1 microscope was used. The images were prepared with Photoshop CS6 software.

### Statistical analysis

Non-parametric 2-way ANOVA test was performed and statistical significance was considered at p<0.05. Statistical analyses of the data were performed using the GraphPad Prism v.8.0 (La Jolla, CA). All the experiments were performed using at least three biological replicates.

## Results and Discussion

### Cre expression from MmuPV1-lox-Cre-lox leads to reporter activation and plasmid recombination *in vitro*

To validate the generated MmuPV1 and control plasmids *in vitro* we transfected mouse embryonic fibroblasts (MEFs) (isolated from R26R-lox-STOP-lox-eYFP mice) with the MmuPV1-lox-Cre-lox plasmid or with the control plasmid, pCAG-Cre. As expected, we detected the function of the Cre recombinase resulting the YFP expression in the transfected cells by either plasmid by immunofluorescence (Figure 1B). Additionally, we isolated DNA from these transfected cells and determined the PCR amplification of the Cre sequence and sequence flanking the Cre cassette (Cre loss), showing the recombination of the MmuPV1-lox-Cre-lox plasmid after Cre expression (Figure 1C).

**Figure 1:**
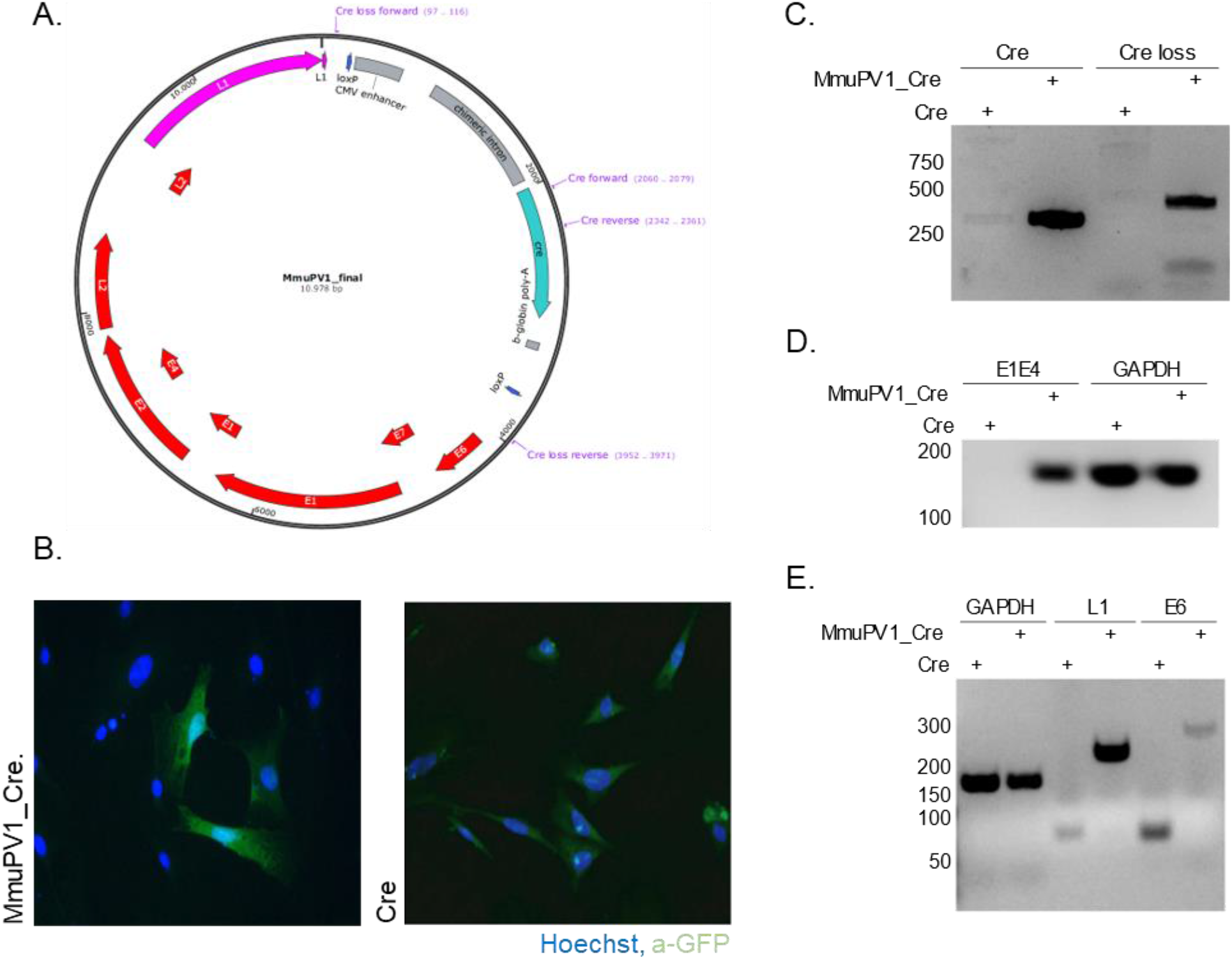
Cre expression from MmuPV1-lox-Cre-lox leads to plasmid recombination *in vitro*. **A)** Map of the MmuPV1-lox-Cre-lox plasmid. **B)** Immunofluorescence of MEFs (isolated from R26R-EYFP mice) transfected with CAG-Cre (Cre) or MmuPV1-lox-Cre-lox (MmuPV1_Cre) plasmid 48 hours post transfection (green: a-GFP, blue: Hoechst). **C)** DNA isolated from transfected cells was used for PCR amplification of the Cre sequence and sequence flanking the Cre cassette to determine recombination of the MmuPV1_Cre plasmid after Cre expression. **D-E)** RT-PCR analysis for the presence of the viral transcripts E1^E4 (E), L1 and E6 (F) in GFP and MmuPV1_Cre transfected MEFs.

Lastly, we detected the presence of MmuPV1 viral transcripts E1-E4, L1 and E6 via RT-PCR analysis in MmuPV1-lox-Cre-lox transfected cells but not in pCAG-Cre transfected cells (Figure 1D-E). Together, these results validate that the constructed MmuPV1-lox-Cre-lox plasmid can indeed lead to reporter activation, Cre-excision, and MmuPV1 genome recircularization *in vitro*.

### Cells which take up MmuPV1-lox-Cre-lox plasmid can be detected and traced over time *in vivo*

Next, we used the plasmid in mice. It is known that infection with MmuPV1 is mainly cleared by the functions of the CD8+ T cell-dependent adaptive immunity (Handisurya et al., 2014). Therefore, only immunocompromised mice are susceptible to the mouse papilloma virus. Thus, we transiently immunocompromised mice before delivering the MmuPV1-lox-cre-lox plasmid by using UVB radiation. UVB was shown to suppress the adaptive immune response whereas the innate immune response remains largely unaffected (Uberoi et al., 2016).

Following the UVB radiation, the base of the tail skin of Rosa26-lox-Stop-lox-YFP mice was superficially abraded with a pipette tip to expose the tail basal layer. Then, MmuPV1-lox-Cre-lox plasmid was delivered and skin samples were harvested at the specified time points to check the presence of YFP expression by immunofluorescence (Figure 2A).

**Figure 2:**
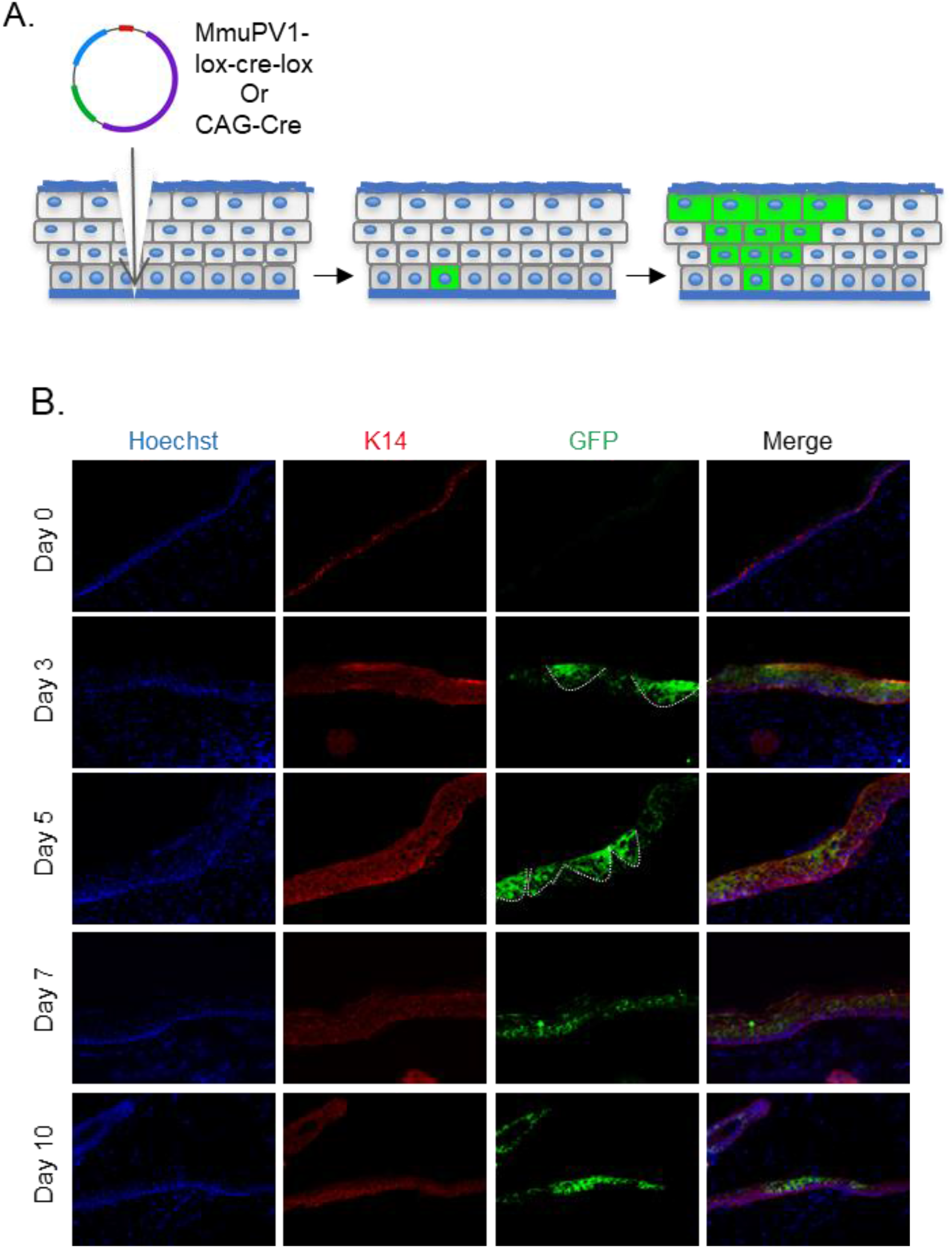
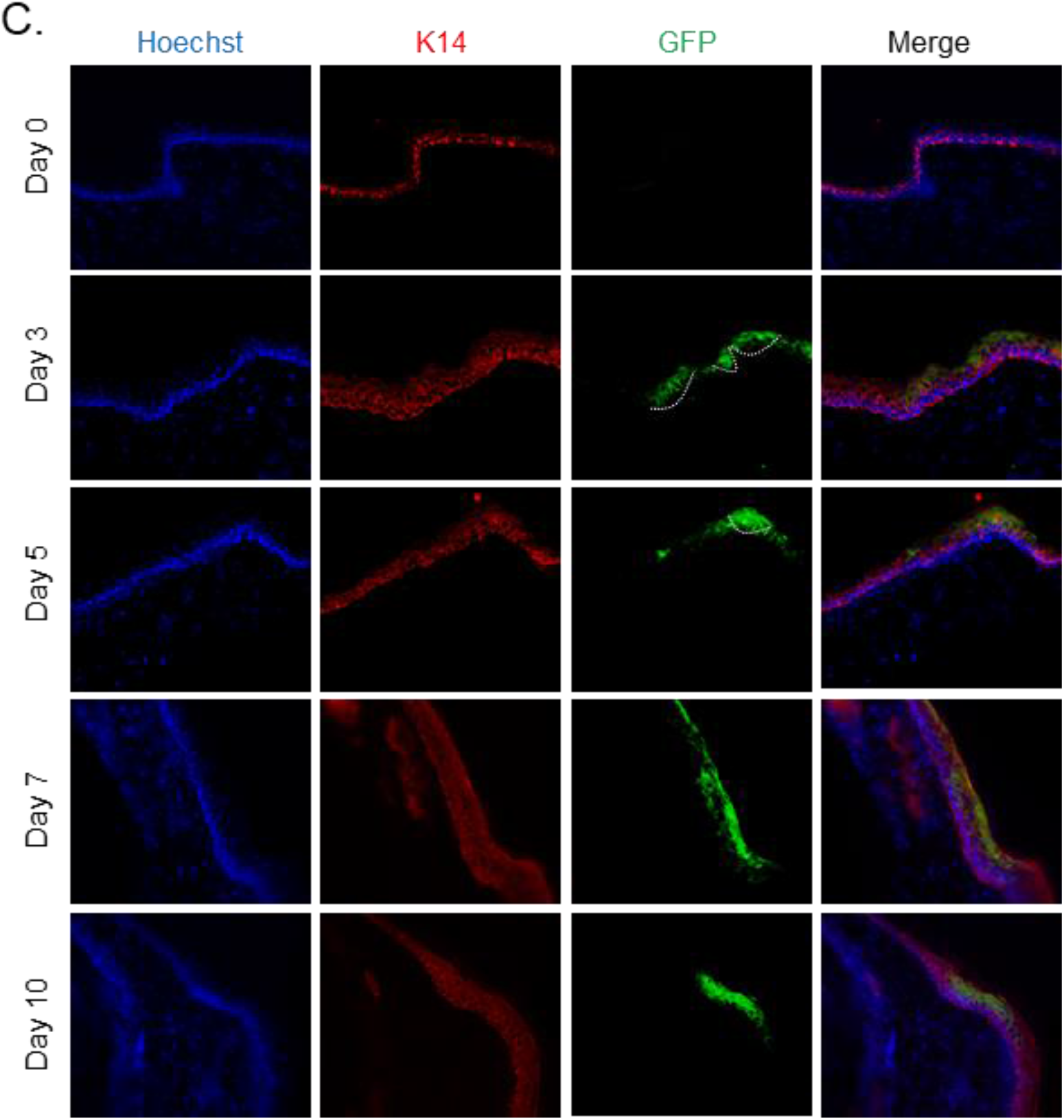
Cells which taken up MmuPV1-lox-Cre-lox plasmid can be detected and traced over time *in vivo*. **A)** Experimental design; The MmuPV1-lox-Cre-lox plasmid is delivered to the tail skin of Rosa26-lox-stop-lox-YFP mice after a UVB irradiation and a superficial scarification. The expected observation is an almost “V” shaped YFP expressing cell population where a single or a couple of basal cells have taken up the plasmid at the bottom and went through cell division where progenies move upwards in the skin layers towards the surface. **B)** Immunofluorescence of MmuPV1-lox-Cre-lox plasmid delivered tail skin section at the indicated time points (red: K14, green: a-GFP). **C)** Immunofluorescence of CAG-Cre plasmid delivered tail skin section at the indicated time points (red: K14, green: a-GFP). White lines indicate the almost “V” shaped YFP expressing cell populations on days 3 and 5 only.

We observed characteristic clones of YFP-expressing cell population in the skin indicative of heritable YFP activation in a skin basal cell. Clones were most prominent at day 3 and day 5 after plasmid delivery (Figure 2B, indicated with white lines). Additionally, we were able to follow-up YFP+ cells at later time points. These results indicate that our model can indeed be used as an *in vivo* lineage tracing model for MmuPV1 where virus-hosting cells and their progeny can be individually observed via their genetically induced YFP expression.

As a control for our lineage tracing model, we use CAG-Cre plasmid in which, drives expression of a self-deleting Cre recombinase, just as in the MmupV1-lox-cre-lox plasmid, but which does not encode any MmuPV1 genes. As performed above, pCAG-Cre was delivered to Rosa26-lox-Stop-lox-YFP mice after UVB irradiation and tail skin scarification. Tails skins were isolated and treated in an identical manner as MmuPV1-treated tissues (Figure 2C, indicated with white lines).

### Tissues treated with MmuPV1-lox-Cre-lox plasmid show evidence of active MmuPV1 gene transcription *in vivo*

We also examined the presence of the viral genes and transcripts in the cells of the same tissue samples and same time points as above. Firstly, we used the RNA in situ hybridization assay of RNAscope to check for the expression of MmuPV1 transcripts, E1-E4 and E6-E7. We observed that in the cells that have taken up the MmupV1-lox-cre-lox plasmid, E6-E7 transcripts started to appear as early as 3 days post-delivery (p.d.) and peaked around day 7 p.d. (Figure 3A) whereas the E1-E4 transcripts were detected at day 5 p.d. onwards and peaked at day 10 p.d. (Figure 3B). Furthermore, we did not detect any of the tested MmuPV1 viral transcripts in the cells that have taken up the CAG-Cre plasmid since the viral genome of MmuPV1 is lacking in this control plasmid (Figure 3C-D).

**Figure 3:**
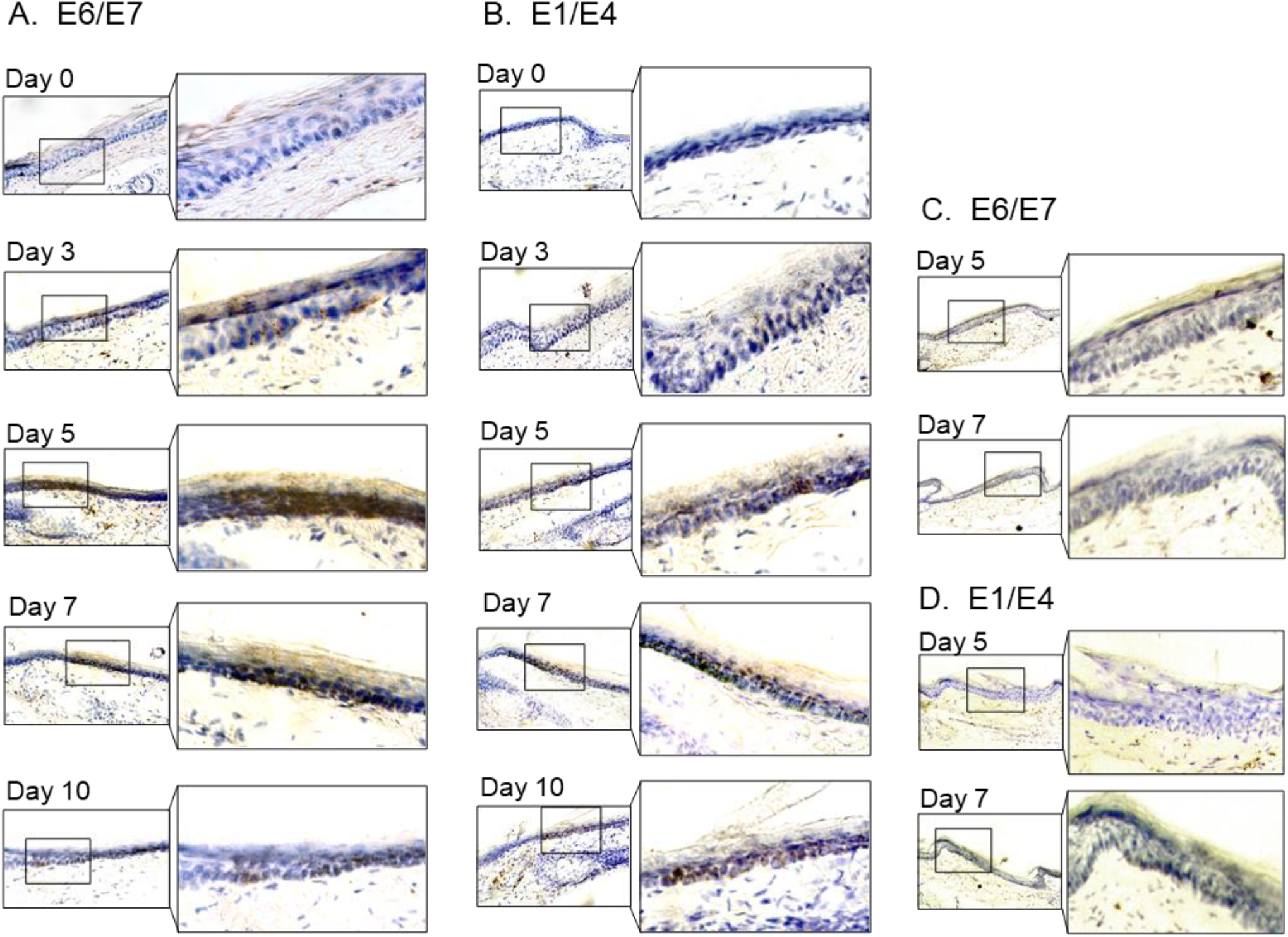
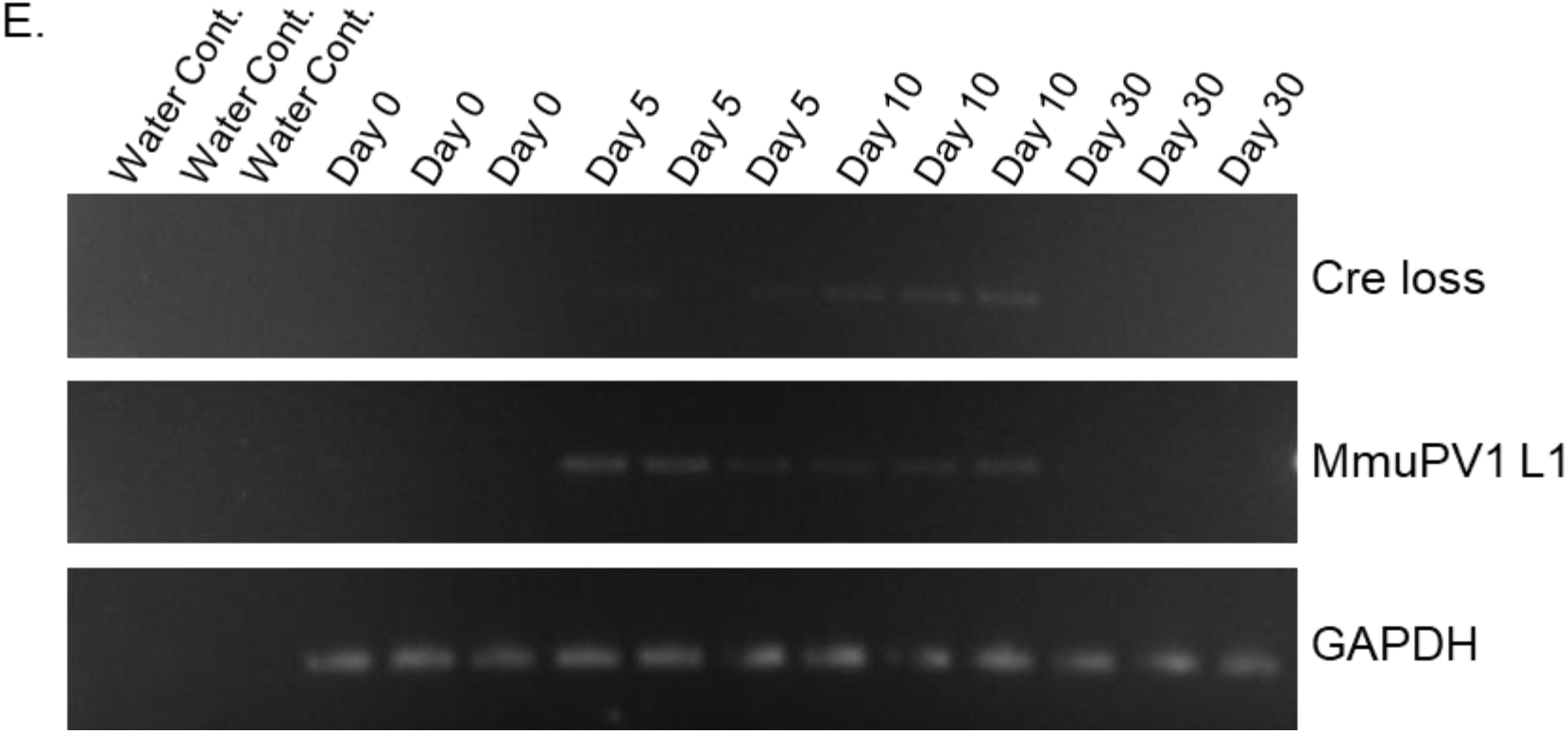
Tissues with MmuPV1-lox-Cre-lox plasmid express viral transcripts *in vivo*. **A-B)** RNA Scope of the MmuPV1-lox-Cre-lox plasmid delivered tail sections at the indicated time points to assess the presence of viral transcripts E6/E7 and E1/E4 respectively. **C-D)** RNA Scope of the CAG-Cre plasmid delivered tail sections at the indicated time points to assess the presence of viral transcripts E6/E7 and E1/E4 respectively. The presence of the transcripts appears as brown dots. **E)** DNA isolated from the swap samples of the MmuPV1-lox-Cre-lox plasmid delivered tail sections and was used for PCR amplification of the sequence flanking the Cre cassette (Cree loss) and L1 of MmuPV1 to determine the presence and the recombination of the MmuPV1_Cre plasmid after Cre expression.

To confirm the presence of genome on the skin surface, we swabbed the surface of the treated tail skins and washed the swaps to remove cells, prior to collecting the skin tissue for the above-mentioned assays. To verify the anticipated recircularization of the MmuPV1 plasmid occurred *in vivo* as well, we isolated DNA to determine the PCR amplification of the sequence flanking the Cre cassette (Cre loss) (Figure 3E). We also assessed the presence of the L1 gene of MmuPV1 via PCR as a way of confirming the presence of the viral genome on the skin surface; GAPDH was used as a housekeeping gene (Figure 3E).

As expected, at day 0 we did not observe any viral transcripts, genes or the Cre loss sequence. However, we detected the Cre loss and the MmuPV1-L1 sequences at day 5 and day 10 samples of the skin surface (Figure 3E). The presence of the re-circularized MmuPV1 plasmid on the surface of the skin could be a result of the eventual migration of the progeny cells to the superficial layer from the initially MmuPV1-harboring cell at the basal layer and shedding the viral particles on the skin surface.

Interestingly, at day 30 time point, we did not detect any viral genome on swabs of the tail skin even though we still detect YFP+ cells at this time point, albeit at low levels (Figure 4A). This might be indicative of the greatly reduced presence (below limit of detection) or clearance of the virus on the skin. Together, we were able to detect the MmuPV1 transcripts of E1-E4 and E6-E7 in the cells which harbor the MmuPV1-lox-cre-lox plasmid and also confirmed the recombination of the MmuPV1-lox-Cre-lox plasmid after Cre expression by detecting the sequence flanking the Cre cassette (Cre loss).

**Figure 4:**
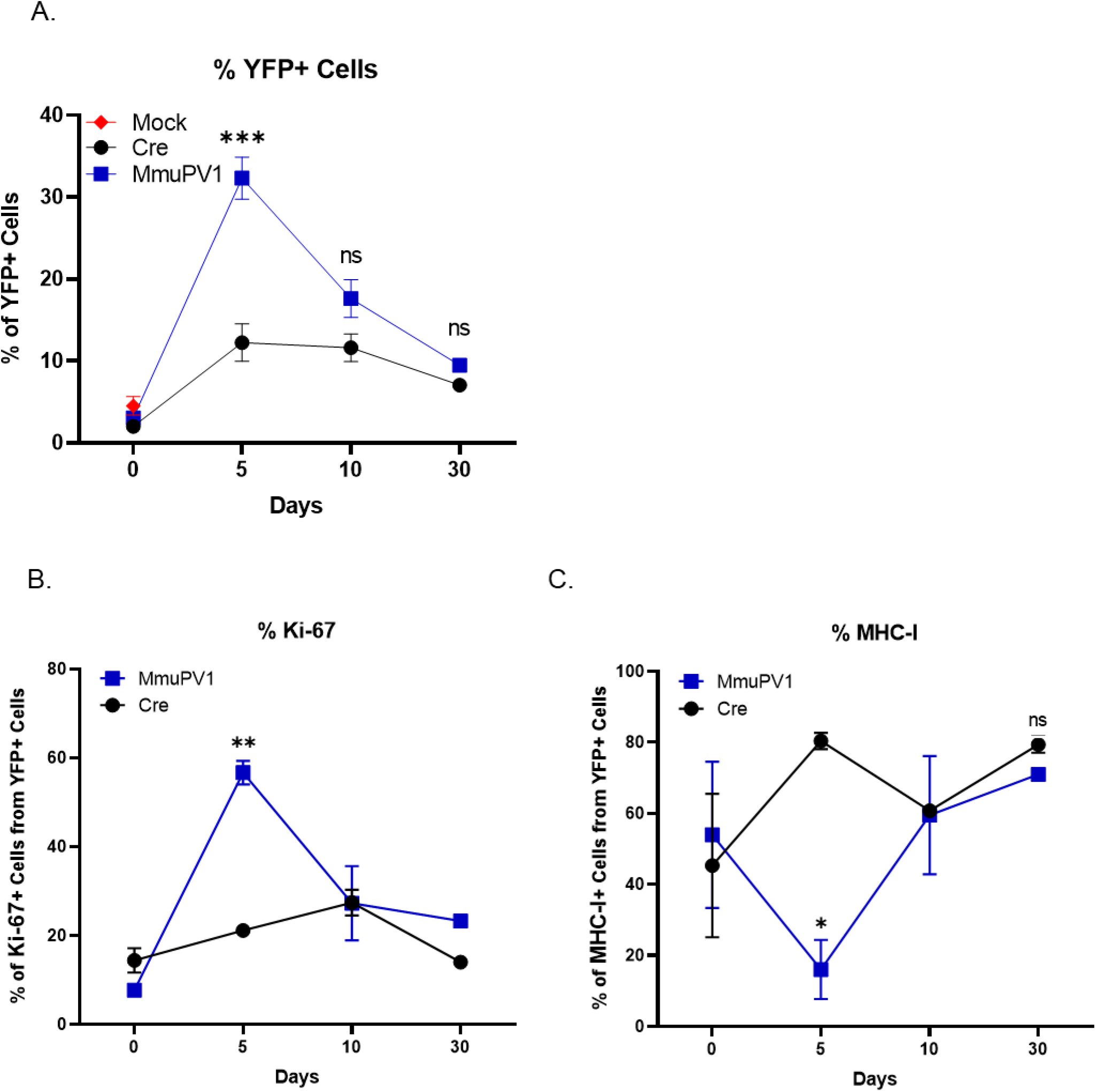
Cells with MmuPV1-lox-Cre-lox plasmid have increased proliferation rate and decreased MHC-I expression on the cell surface. **A)** Percentages of YFP+ cells. **B)** Percentages of Ki-67+ cells out of only YFP+ cells. **C)** Percentages of MHC-I+ cells out of only YFP+ cells. Mock infection with PBS (red square) was only performed for day 0 time point. Data representative of two independent experiments with n=3 mice per group. The results are presented as means ± SEM. not significant (ns), * p<0.05, ** p<0.01 and *** p<0.001 by nonparametric 2-way ANOVA test.

### Progeny of cells which received MmuPV1-lox-Cre-lox plasmid exhibit higher proliferation dynamics and lower surface expression of MHC-I *in vivo*

To provide proof of principle that our model system may be used to answer question regarding papillomavirus biology, and taking advantage of the ability to detect YFP expressing cells, we applied the plasmids to the tail skin of Rosa26-lox-Stop-lox-YFP mice as mentioned above. Then, skin samples were harvested at the specified time points and processed and stained for flow cytometric analysis.

Firstly, we applied a standard gating strategy to exclude dead and doublet cell populations (Figure S1A). Then, we detected the YFP expressing cells and compared numbers and frequencies between groups and different time points (Figure 4A, Figure S1B). We found that both numbers and percentages of YFP+ cells were much higher in MmuPV1 samples at day 5 p.d., which decreased overtime as seen at day 10 and day 30 p.d. Whereas, pCAG-Cre samples only start to have increased YFP+ cells around day 10 p.d. (Figure 4A, Figure S1B). These results are also in line with the immunofluorescence data we saw previously and can indicate to a difference in the actual amount of plasmid delivered to the tissue or the different dynamics of the plasmids due to the presence of the MmuPV1 genes. Furthermore, coinciding with the increased number of YFP+ cells, we observed that MmuPV1-harboring cells exhibit significantly increased levels of Ki-67 around day 5 p.d. compare to other time points (Figure 4B, Figure S1C).

Papillomaviruses have previously been reported to reduce MHC-I expression, but evidence was generated *in vitro* (Georgopoulos et al., 2000) (Heller et al., 2011) (Li et al., 2010). To evaluate the levels of MHC-I specifically on the cell surface of the MmuPV1-harboring cells, we used flow-cytometry to interrogate MHC-I levels within the YFP+ population. Interestingly, we observed significantly lower expression of MHC-I molecules on the YFP+ cells of MmuPV1-treated tissues at day 5 p.d. whereas YFP+ cells in control treated tissues exhibit increased MHC-I expression at this time point, presumable due to YFP neo-antigen expression (Figure 4C, Figure S1D). These effects waned at later time points. Our results agree with previous *in vitro* studies and indicate a possible immune evasion strategy by the MmuPV1 where the virus is causing a decreased expression of MHC-I molecules on the cell surface to reduce the antigen exposure to the immune cells.

It is important to note, that the differences in proliferation and MHC- I expression were specific to the YFP+ cells within the tissue. Also, within the same tissues, we analyzed the Ki-67 and MHC-I levels of the neighboring YFP- cells which haven’t taken up the given plasmid (Figure S2). We observed no significant difference between time points, or between the YFP- cell population of the MmuPV1 plasmid-treated tissue and the CAG-Cre plasmid-treated tissue (Figure S2A-B). These results suggest that any differences observed are specific to plasmid uptake.

Together, these results indicate that during early acute infection MmuPV1 causes increased proliferation which may be accompanied by decreased antigen presentation. Additionally, here we show that our novel *in vivo* lineage tracing model can be used as a tool to answer such key questions in the field of MmuPV1 biology.

## Conclusions and significance

We describe here a novel *in vivo* lineage tracing model for MmuPV1 infected cells in laboratory mice (*Mus musculus*). We propose that the ability to trace MmuPV-1 harboring cells and their progeny by microscopy, as well as to quantify and analyze by means of flow-cytometry renders this model a suitable and convenient tool for answering a variety of outstanding questions in papillomavirus biology (Lambert et al., 2020).

---

**Supplementary Figure 1:**
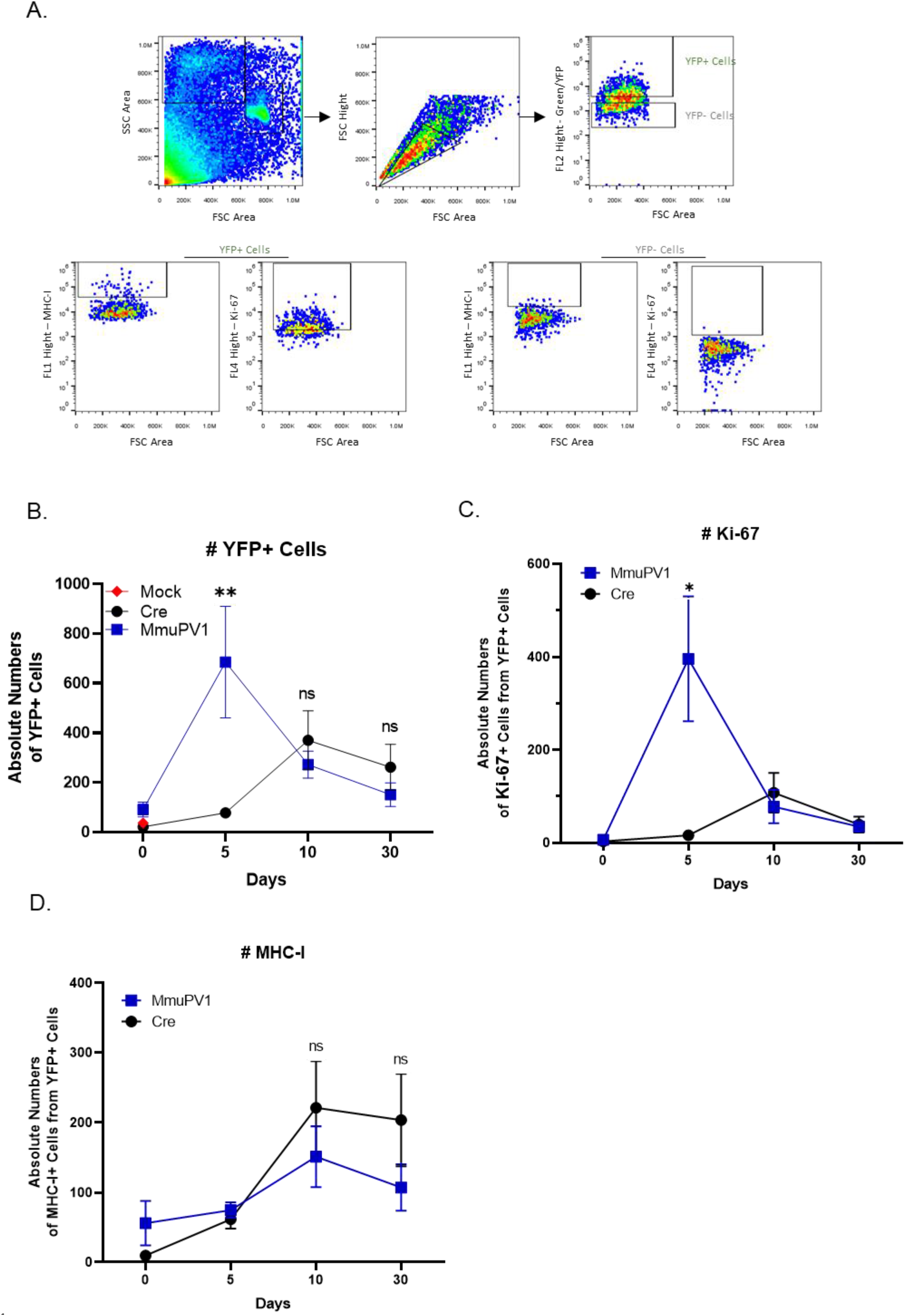
Cells with either MmuPV1-lox-Cre-lox or control plasmid can be detected and analyzed by flow cytometry. **A)** The population of cells was detected depending on their size and complexity, omitting the dead cells (FSC-SSC gate) and later excluding the doublets (FSC-Height/FSC-Area gate). **B)** Absolute numbers of YFP+ cells. **C)** Absolute numbers of Ki-67+ cells out of only YFP+ cells. **D)** Absolute numbers of MHC-I+ cells out of only YFP+ cells. Data representative of two independent experiments with n=3 mice per group. The results are presented as means ± SEM. not significant (ns), * p<0.05, ** p<0.01 and *** p<0.001 by nonparametric 2-way ANOVA test.

**Supplementary Figure 2:**
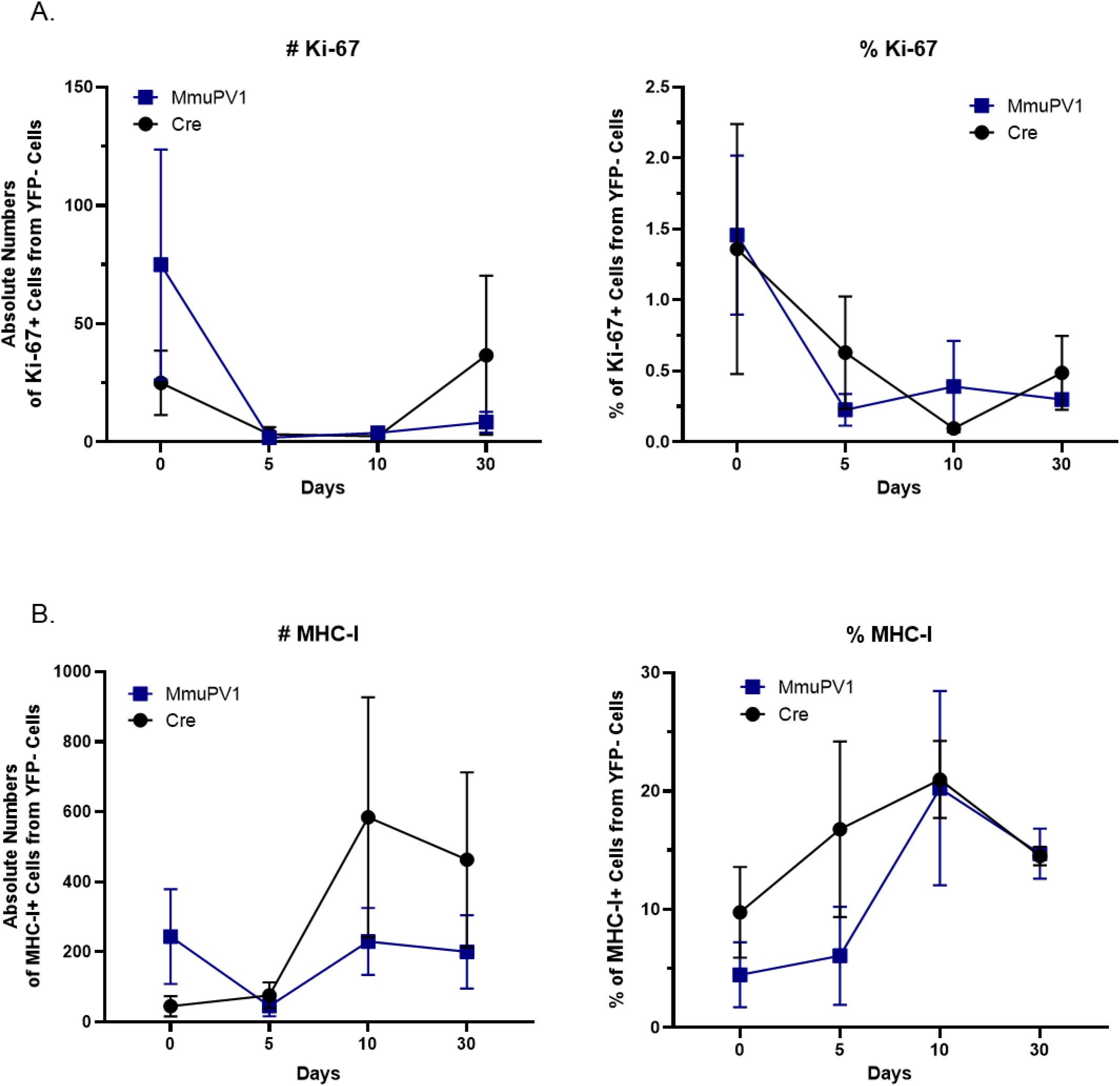
YFP- Cells in the tissues treated with MmuPV1-lox-Cre-lox and CAG-Cre plasmid have similar Ki-67 and MHC-I expression. **A)** Absolute numbers (left panel) and percentages (right panel) of Ki-67+ cells out of only YFP-cells. **B)** Absolute numbers (left panel) and percentages (right panel) of MHC-I+ cells out of only YFP- cells. Data representative of two independent experiments with n=3 mice per group. The results are presented as means ± SEM. not significant (ns), * p<0.05, ** p<0.01 and *** p<0.001 by nonparametric 2-way ANOVA test.

